# A single-administration therapeutic interfering particle reduces SARS-CoV-2 viral shedding and pathogenesis in hamsters

**DOI:** 10.1101/2022.08.10.503534

**Authors:** Sonali Chaturvedi, Nathan Beutler, Michael Pablo, Gustavo Vasen, Xinyue Chen, Giuliana Calia, Lauren Buie, Robert Rodick, Davey Smith, Thomas Rogers, Leor S. Weinberger

## Abstract

The high transmissibility of SARS-CoV-2 is a primary driver of the COVID-19 pandemic. While existing interventions prevent severe disease, they exhibit mixed efficacy in preventing transmission, presumably due to their limited antiviral effects in the respiratory mucosa, whereas interventions targeting the sites of viral replication might more effectively limit respiratory virus transmission. Recently, intranasally administered RNA-based therapeutic interfering particles (TIPs) were reported to suppress SARS-CoV-2 replication, exhibit a high barrier to resistance, and prevent serious disease in hamsters. Since TIPs intrinsically target the tissues with the highest viral replication burden (i.e., respiratory tissues for SARS-CoV-2), we tested the potential of TIP intervention to reduce SARS-CoV-2 shedding. Here, we report that a single, post-exposure TIP dose lowers SARS-CoV-2 nasal shedding and at 5 days post-infection infectious virus shed is below detection limits in 4 out of 5 infected animals. Furthermore, TIPs reduce shedding of Delta variant or WA-1 from infected to uninfected hamsters. Co-housed ‘contact’ animals exposed to infected, TIP-treated, animals exhibited significantly lower viral loads, reduced inflammatory cytokines, no severe lung pathology, and shortened shedding duration compared to animals co-housed with untreated infected animals. TIPs may represent an effective countermeasure to limit SARS-CoV-2 transmission.

**Significance:** COVID-19 vaccines are exceptionally effective in preventing severe disease and death, but they have mixed efficacy in preventing virus transmission, consistent with established literature that parenteral vaccines for other viruses fail to prevent mucosal virus shedding or transmission. Likewise, small-molecule antivirals, while effective in reducing viral-disease pathogenesis, also appear to have inconsistent efficacy in preventing respiratory virus transmission including for SARS-CoV-2. Recently, we reported the discovery of a single-administration antiviral Therapeutic Interfering Particle (TIP) against SARS-CoV-2 that prevents severe disease in hamsters and exhibits a high genetic barrier to the evolution of resistance. Here, we report that TIP intervention also reduces SARS-CoV-2 transmission between hamsters.

## Introduction

Interrupting transmission of respiratory viruses remains a fundamental medical and public-health challenge. While COVID-19 vaccines are exceptionally effective in preventing severe disease and death, accumulating data show they have mixed efficacy in preventing viral transmission (1-5), consistent with established literature that parenteral vaccines for other viruses fail to prevent mucosal virus shedding or transmission (6-8). Small-molecule antivirals, while effective in reducing viral-disease pathogenesis, also appear to have inconsistent efficacy in preventing respiratory virus transmission (9, 10), including for SARS-CoV-2 (11-13), possibly due to slow diffusion into the respiratory tract (14). Antibody-based treatments for SARS-CoV-2, which are susceptible to escape (15, 16), appear similarly unable to limit virus shedding or transmission (17), consistent with previous challenges in preventing acute respiratory infections (18). Historically, it has been proposed that interventions targeting the sites of viral replication might more effectively limit respiratory virus transmission (19, 20), but this has been challenging to demonstrate experimentally and has not been achieved in practice.

Recently, we reported that a single dose of an intranasally administered mRNA-based therapeutic interfering particle (TIP) substantially reduces SARS-CoV-2 replication, pathogenesis, and disease in Syrian golden hamsters, and exhibits a high genetic barrier to the evolution of resistance (21). Based upon the historical phenomenon of defective interfering particles (DIPs) (22-24), TIPs encode only a small, non-coding portion of the viral genome (< 2 kb in the case of SARS-CoV-2 (25)) and lack self-replication but, distinct from DIPs, conditionally replicate with a reproductive number > 1 (26, 27). As obligate intracellular molecular parasites, TIPs suppress viral burst size and reduce cell-to-cell virus transmission, thereby limiting disease pathogenesis (21). The molecular mechanism of interference and conditional R_0_>1 replication intrinsically target the TIP antiviral effect to the tissues with the highest viral replication burden (i.e., respiratory tissues for SARS-CoV-2). Therefore, we tested the potential of TIP intervention to reduce SARS-CoV-2 viral transmission (**Fig. 1a)**.

**Figure 1:**
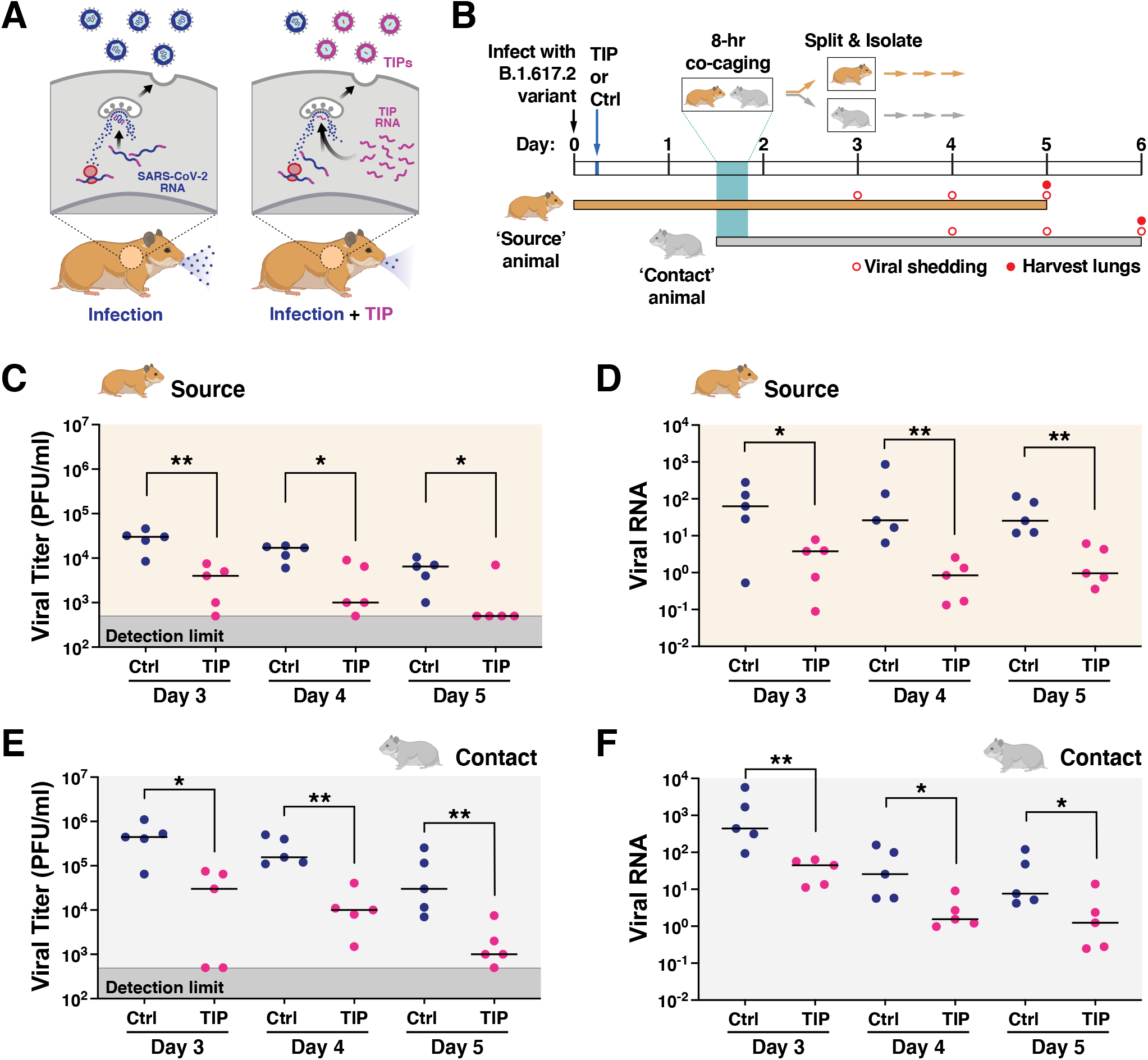
TIPs reduce nasal shedding of highly pathogenic SARS-CoV-2 (B.1.617.2) in Syrian golden hamsters. (**A)** Schematic of putative mechanism-of-action for TIP-mediated reduction in virus transmission. **(B)** Schematic of experimental design. Syrian golden hamsters (source) were intranasally infected with 10^6^ plaque forming units (PFU) of SARS-CoV-2 (B.1.617.2). At 6 h post-infection, TIP RNA LNPs (n=5) or Ctrl RNA LNPs (n=5) were intranasally administered to source animals via instillation, and hamsters were caged alone until 36 h post-infection. At 36 h, an uninfected untreated (contact) hamster was co-caged with each source hamster for 8 hours. At 44 h post-infection, source and contact hamsters were separated and caged alone for the duration of the study. Nasal washes were performed at day 3 (source only), day 4 (source and contact), day 5 (source and contact), and day 6 (contact only). Source hamsters were sacrificed at day 5, and contact hamsters sacrificed at day 6, to harvest lungs. **(C)** TIP treatment reduces infectious virus shedding in nasal washes. Infectious SARS-CoV-2 in nasal washes (for day 3, 4 and 5) of source animals treated with TIP (n=5) or Ctrl LNPs (n=5), quantified by plaque assay. **(D)** TIP treatment reduces SARS-CoV-2 RNA shedding in nasal washes. Viral RNA was extracted from the nasal washes for TIP (n=5) or Ctrl (n=5) treated source animals at day 3, 4 and 5, and quantified by qRT-PCR for N gene. **(E)** Contacts of TIP-treated animals exhibit reduced infectious virus in nasal washes. Quantification of infectious SARS-CoV-2 from the nasal washes (for day 4, 5 and 6) for contacts of TIP (n=5) or Ctrl (n=5) treated animals was performed using plaque assay. **(F)** Contacts of TIP-treated animals exhibit reduced viral RNA in nasal washes. Viral RNA was extracted from the nasal washes from contacts of TIP-or Ctrl-treated animals at day 4, 5 and 6, and quantified by qRT-PCR for N gene. Medians of each arm are shown as black horizontal bars. For all panels: **, p < 0.01; *, p < 0.05 from Mann-Whitney U test.

## Results & Discussion

### Design of Co-caging Transmission Studies in Syrian Golden Hamsters

To test if TIPs reduce SARS-CoV-2 transmission, we employed the Syrian golden hamster model of infection (28) based on a previously reported experimental scheme (**Fig. 1b**). Briefly, a group of Syrian golden hamsters were intranasally inoculated with 10^6^ plaque forming units (PFU) of SARS-CoV-2 B.1.617.2 (a.k.a., Delta variant) and designated as ‘source’ animals. At 6 hours post-infection, source animals received a single intranasal dose of either TIP RNA lipid nanoparticles (LNPs) or Ctrl RNA LNPs (n=5 per group). When the source animals were near peak infectivity (36 hours post-infection), each animal was co-caged for 8 hours with an uninfected, untreated hamster (i.e., ‘contact’ animal), to promote efficient (aerosol and fomite) transmission of SARS-CoV-2. At 44 hours post-infection, the source and contact animals were separated into individual cages for the duration of the study. Nasal washes were then collected daily, and source and contact animals were sacrificed on day 5 and 6, respectively (i.e., ∼5 days post infection for each), to harvest lungs for viral titering and analysis of histopathology and inflammation.

### TIPs reduce transmission of highly pathogenic SARS-CoV-2 (B.1.617.2) in Syrian golden hamsters

TIP-treated ‘source’ hamsters exhibited significantly lower virus shedding in daily nasal washes, as measured by infectious virus (**Fig. 1c**) or viral RNA (**Fig. 1d**), and exhibited faster decays in nasal viral loads. Strikingly, infectious virus shed from TIP-treated animals decayed to below the limit of detection (LOD) by day 5 post infection in four out of five animals, whereas all Ctrl-treated source animals shed high levels of virus to day 5 **(*SI Appendix*, Fig. S1)**. Similar reductions in nasal virus shedding were also observed for contacts of TIP-vs. Ctrl-treated animals **(Fig. 1e, f)**.

In both source and contact animals, lungs were harvested at 5- and 6-day post infection (**Fig. 2a**) and virus titer quantification was performed. TIP treatment reduced infectious SARS-CoV-2 viral load in the lungs by >3 Logs (**Fig. 2b**), and reduced SARS-CoV-2 RNA levels by >2 Logs **(Fig. 2c)**. Consistent with previous reports (28, 29), contact hamsters exhibited higher viral burden than source hamsters, possibly due to the mode of inoculation—the comparatively extended 8-hour aerosol plus fomite inoculation—resulting in a higher viral inoculum and/or delayed viral dynamics in the contact animals.

**Figure 2:**
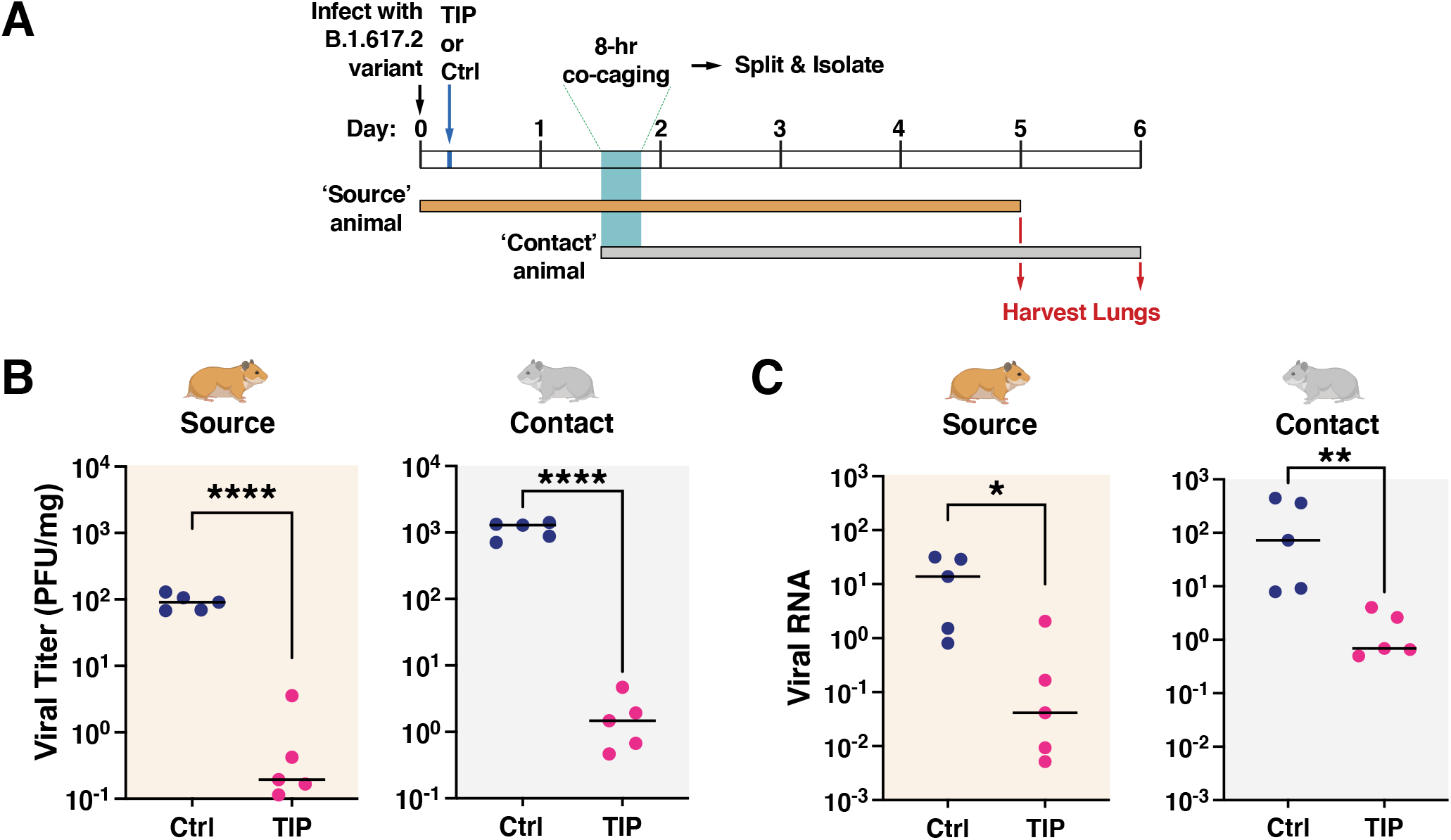
TIPs reduce transmission of SARS-CoV-2 (B.1.617.2) in the lungs of Syrian golden hamsters. **(A)** TIP treatment reduces infectious viral load in lungs of source hamsters and their contacts. Quantification of infectious SARS-CoV-2 in lungs of source and contact animals (n=5 for all arms) as analyzed by plaque assay. **(B)** TIP treatment reduces viral RNA in lungs of source hamsters and their contacts. Total RNA in lungs was harvested from source and contact animals quantified by qRT-PCR using primers specific for N gene of SARS-CoV-2 and normalized to beta-actin. Medians of each arm are shown as black horizontal bars. For all panels: ****, p<0.0001, **, p < 0.01; *, p < 0.05 from Mann-Whitney U test.

### TIP-mediated reduction in SARS-CoV-2 transmission reduces disease pathogenicity in both source and contact animals

Analysis of inflammatory gene expression in animal lungs showed significant reductions in inflammation in TIP-treated source animals, consistent with our previous findings (21), and significant reductions in inflammation in the contacts of TIP-treated animals (**Fig. 3a**). Histopathological analysis also showed substantial improvement in lung disease of TIP-treated animals, also consistent with previous data (21), as well as substantial improvement in lung disease of the contacts of TIP-treated animals (**Fig. 3b–c; *SI Appendix*, Fig. S2**). Since host inflammation and lung damage reflect time-integrated viral burden—whereas viral load measurements are a temporal snapshot—these inflammatory and histopathological data support the hypothesis that TIP intervention reduces viral shedding and transmission of SARS-CoV-2, and do not support the alternate hypothesis that viral load in source animals peaked at a time not captured by the nasal-wash collection schedule. Moreover, the increased viral load in the contact animals indicates that pathogenesis was not simply delayed in the contact animals compared to the source animals. Overall, the data indicate that a single intranasal dose of TIP LNPs reduces shedding and transmission of SARS-CoV-2, thus conferring protection to contact animals.

**Figure 3:**
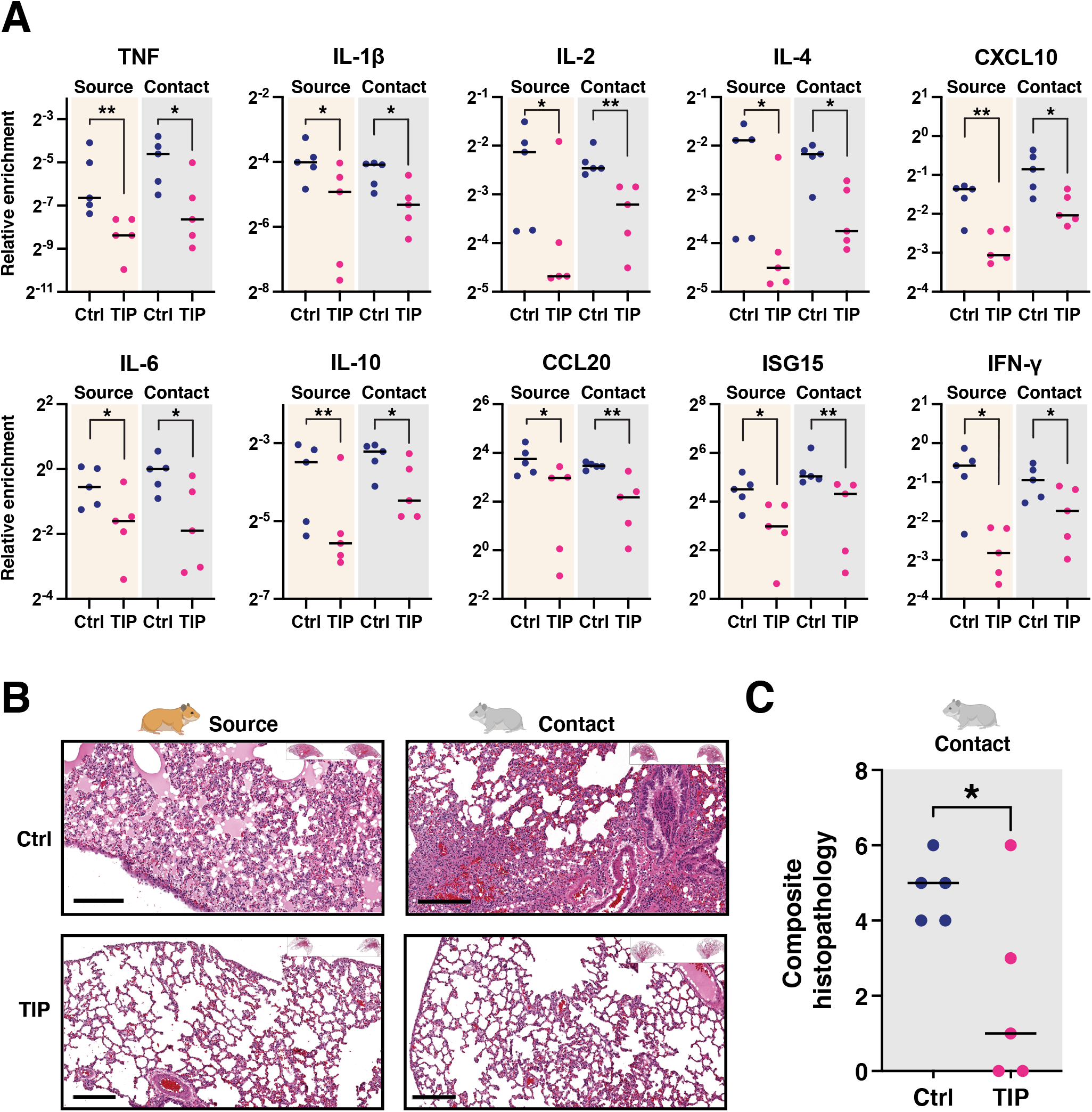
TIP-mediated reduction in SARS-CoV-2 transmission reduces disease pathogenicity in both source and contact animals. **(A)** qRT-PCR expression analysis for pro-inflammatory and interferon-stimulated genes in hamster lungs. Lungs were harvested from TIP-treated (n=5) or Ctrl-treated (n=5) source animals (at day 5) and their respective contact hamsters (at day 6), homogenized, and analyzed by qRT-PCR for pro-inflammatory cytokines and interferon-stimulated genes using respective primers and normalized to beta-actin. **, p < 0.01; *, p < 0.05 from Mann-Whitney U test. (**B**) H&E staining of representative lung sections of TIP- and Ctrl-treated source animals and their contacts. Scale bars=200 *μ*m. **(C)** Histopathological scoring of lung sections for the prevalence of pulmonary infiltrates, edema, macrophages, and septum widening, resulting in a composite histopathological score ranging from a minimum of 0 (indicating the absence of visual indications of pathogenicity in all scoring dimensions), to a maximum of 8 (indicating end-stage pathogenesis evidenced by overwhelming infiltrates, and/or edema, macrophage, and septum widening). Medians of each arm are shown as black horizontal bars. *, p < 0.05 from Student’s t test.

To test a second alternate hypothesis that TIPs might be mobilizing from source animals to therapeutically interfere and lower viral load within the contact animal, we used qRT-PCR to assay TIP RNA in contact animals and found no evidence of TIP transmission (***SI Appendix*, Fig. S3**). These data are consistent with our previous model analysis (21) predicting that, while cell-to-cell transmission is efficient in the case of SARS-CoV-2, between-host TIP transmission faces several bottlenecks resulting in an R_0_ << 1.

### TIP-mediated transmission reduction is robust across SARS-CoV-2 strains

To verify that these transmission results were not limited to the Delta variant (B.1.617.2), we also conducted an analogous hamster transmission experiment, with the same TIP RNA, using the archival SARS-CoV-2 WA-1 strain and observed qualitatively similar results (***SI Appendix*, Fig. S4**), indicating that TIP treatment reduces viral shedding, pathogenesis, and transmission across multiple viral strains.

### Viral dynamics models reveal TIP-mediated reductions in viral shedding in contact animals

Next, to determine if contacts of TIP-treated animals also showed reduced viral shedding, we extended an established mathematical model of viral dynamics (30) to include TIPs and then forecast the duration of infectious virus shedding from contact-animal nasal-wash data. Since we observed no evidence of transmission of TIPs, the viral dynamics in contact animal were modeled without any TIPs (**Fig. 4a**). The model was first benchmarked against data from each individual source animal (**Fig. 4b**), and it estimated that TIP-treated source animals stopped shedding 2 days faster than Ctrl-treated animals (**Fig. 4d**), in agreement with our experimental data (**Fig. 1c; *SI Appendix*, Fig. S1**). The model also indicated that TIP treatment in source animals generated a ∼1-Log reduction in the peak level of virus shed and in the total amount of virus shed during co-housing (**Fig. 4e, f**). Next, changes in contact animal shedding were estimated by fitting the now-benchmarked model to the nasal wash viral titer data from each contact hamster (**Fig. 4c**). The model indicated that contacts of TIP-treated animals stop shedding 2 days faster than the contacts of Ctrl-treated animals (**Fig. 4g**). The contacts of TIP-treated animals also exhibit a ∼1-Log reduction in peak shedding compared to the contacts of Ctrl-treated animals (**Fig. 4h**). Collectively, this analysis suggests that post-exposure TIP treatment of infected hamsters lowers transmission by reducing both the level and duration of virus shed in both source and contact animals.

**Figure 4:**
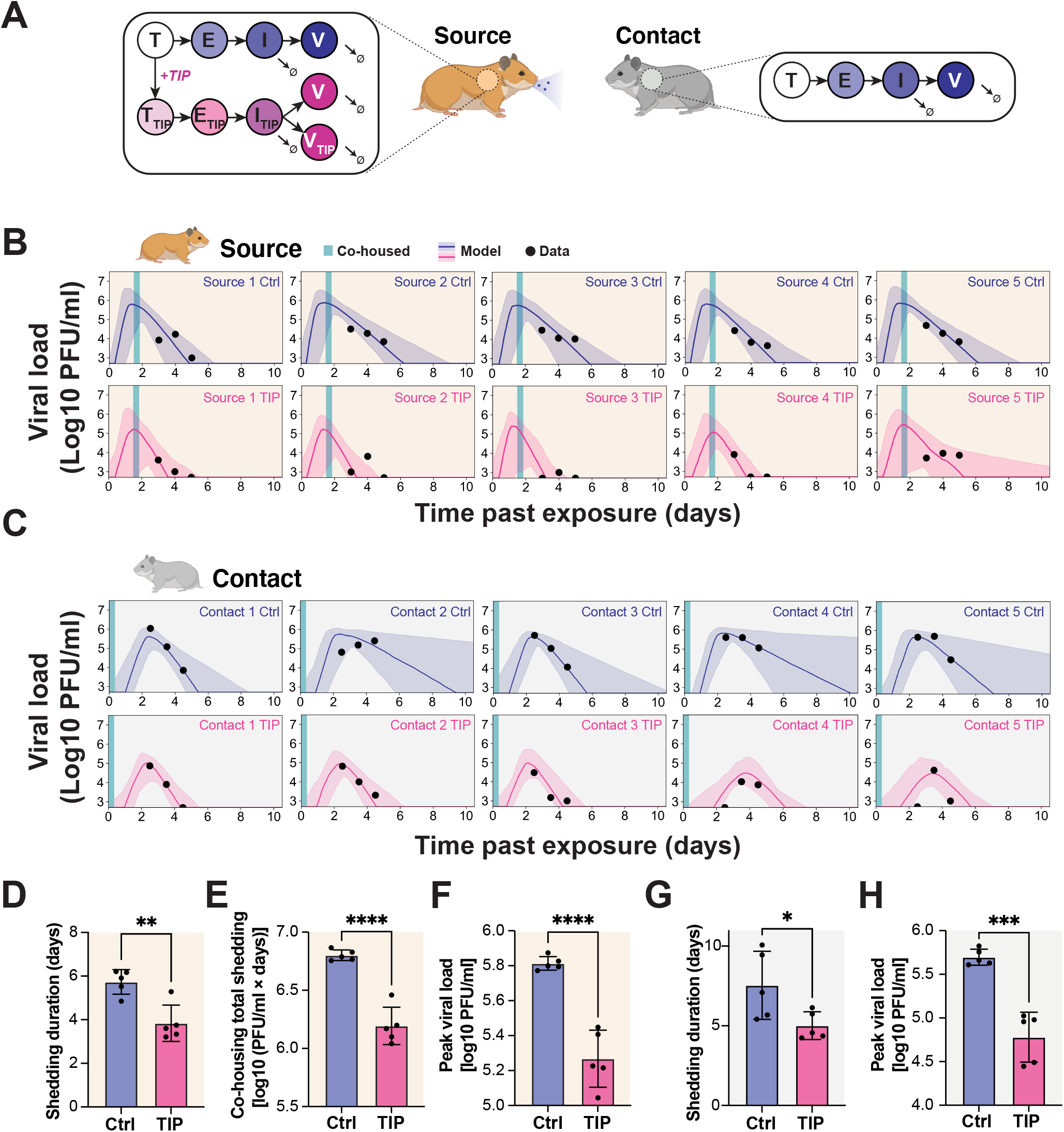
Viral dynamics models reveal TIP-mediated reductions in viral shedding in contact animals. **(A)** Schematic of *in silico* model for SARS-CoV-2 dynamics in source and contact hamsters. **(B)** Viral dynamics models fit to nasal wash plaque assay data from source hamsters. A 95% credible interval surrounding a median prediction line is shown. The abscissa uses the time past exposure, which for source hamsters begins at t=0 hours post-infection. **(C)** Viral dynamics models fit to nasal wash plaque assay data from contact hamsters. A 95% credible interval surrounding a median prediction line is shown. The abscissa uses the time past exposure, which for contact hamsters begins at t=36 hours post infection of source hamsters. **(D)** Model inference shows that TIP reduces the duration of viral shedding in source hamsters, defined as the time until the simulated viral load dropped below the experimental limit of detection. **(E)** Model inference shows that TIP reduces the total viral shedding in source hamsters during the co-housing period, defined as the integral of the simulated viral load during co-housing. (**F**) Model inference shows that TIP reduces the peak viral shedding in source hamsters. (**G**) Model inference shows that contacts of TIP-treated animals exhibit reduced duration of viral shedding. (**H**) Model inference shows that contacts of TIP-treated animals exhibit reduced peak viral load. For panels (**D-H**), each dot represents a model fit to the time course of an individual hamster; TIP (n=5), Ctrl (n=5). ****, p<0.0001; ***, p<0.001; **, p<0.01; *, p<0.05 from Student’s t test.

Overall, these data provide proof-of-concept that a single-administration, post-exposure intervention using mRNA-based TIP LNPs reduces the amount and duration of SARS-CoV-2 virus shedding. The data indicate that TIPs are effective against diverse archival variants (i.e., WA-1) as well as more recent highly pathogenic variants of concerns (e.g., Delta variant).

Our study has several limitations. First and foremost, the intervention was unable to fully eliminate virus transmission from source animals, since contact animals did become infected (i.e., TIP intervention did not generate transmission ‘sterilization’). However, as noted, similar experimental designs (9-14, 17) were employed for other antiviral interventions and failed to generate any reduction in virus transmission in hamsters. It is also possible that our experimental design of 8 hours of co-housing, allowing both aerosol and fomite transmission, resulted in highly efficient, super-physiological, transmission that is not reflective of what might occur between humans.

Recently, a pre-exposure prophylaxis intervention for an oral antiviral protease inhibitor was reported to inhibit transmission between hamsters (31), albeit with a substantially lower virus inoculation of 10^4^ PFU than used in this study (10^6^ PFU in this study) and a reduced study duration (3–4 day duration vs. 5–6 day duration in this study). Regardless, it is possible that pre-exposure TIP prophylaxis may similarly result in more effective transmission reduction given our previous results (21).

Notably, translating these results to humans will require further study given the significant physiological differences between primates and rodents (e.g., in nasal turbinate architecture and the rate of disease progression). The disease course in SARS-CoV-2 infected hamsters is highly accelerated compared to humans, such that 6-12 hrs post infection corresponds to ∼1.25–3.5 days, in humans. Specifically, hamsters typically clear infectious virus (not RNA) by day 4 after peak load (28), whereas humans appear infectious 20–30 days post peak load (32-34). Consequently, there appears to be roughly a 5–8x accelerated disease course in hamsters compared to humans. This timing may be comparable to that used for other SARS-CoV-2 antivirals that show therapeutic efficacy if administered within the first few days after onset of symptoms.

The computational models we employed also face the common limitations of such models, and the predictions will require further testing of transmission to secondary and tertiary contacts. However, the results of such secondary/tertiary transmission studies will be highly sensitive to experimental design, and will require extensive testing of alternate timing and duration of the co-housing scenarios. Whereas the current TIPs do not appear to efficiently transmit between hosts due to transmission bottlenecks, models predict (26, 27) that TIPs could either be engineered to transmit and thus improve population-level efficacy of the intervention, or be engineered to further prevent host transmission as a safety measure. Broadly, the data herein validate the concept that antiviral interventions which specifically target the site of viral replication may effectively reduce respiratory virus transmission.

## Abbreviations

(TIP): therapeutic interfering particle
(LNP): lipid nanoparticle
(LOD): limit of detection
(R_0_): basic reproductive ration

## Acknowledgements

We thank K. Claiborn for editing, Blaise Ndjamen and F.N.N. Pitchai for technical guidance; and the Gladstone-UCSF CFAR flow-cytometry core, funded by NIH grants P30 AI027763 and S10 RR028962 and the James B. Pendleton Trust, as well as the Gladstone Genomics and Histology Cores.

## Funding

This work was supported by Pamela and Edward Taft (to L.S.W.), US Army Medical Infectious Disease Research Program (MTEC 2020-492) (to L.S.W.), and NIH DP1DA051144 (to L.S.W.).

## Author contributions

S.C. and L.S.W conceived and designed the study, N.B. performed hamster experiments, M.P. conceived, designed and performed modelling for the study, S.C., G.V. and G.C. designed, performed and analysed the lipid-nanoparticle studies, S.C., N.B., M.P., G.V., X.C., G.C., L.B., R.R., D.S., T.R., designed and performed the experiments and curated the data. R.R. and L.S.W provided reagents and resources. S.C., M.P., G.V., R.R., D.S., and L.S.W. wrote the paper.

## Declaration of Interests

L.S.W., S.C., and R.R. are co-inventors on a patent application filed for therapeutic interfering particles for SARS-CoV-2. L.S.W. is a scientific co-founder of VxBiosciences.

## Data and materials availability

All data, code, and materials used in the analysis are available in the manuscript and Zenodo (https://doi.org/10.5281/zenodo.6762604).

## Materials and Methods

### Virus and cell culture conditions

SARS-CoV-2 isolate (USA-WA1/2020) and SARS-CoV-2 variant (B.1.617.2) were obtained from Biodefense and Emerging Infections (BEI) Resources. Vero E6 cells were used to prepare viral stocks in Dulbecco’s modified Eagle’s medium (DMEM) supplemented with 10% fetal bovine serum (FBS) and 1% penicillin and streptomycin (P/S). All live virus experiments were performed at the Gladstone Institutes in a Biosafety Level 3 (BSL3) containment facility, or at the Scripps Research Institute in an Animal Biosafety Level 3 (ABSL3) containment facility. All live virus experiments at Gladstone were performed under an approved Biosafety Use Authorization from University of California San Francisco (UCSF) in compliance with institutional guidelines and procedures. All live virus experiments at Scripps were performed under an approved Biosafety Use Authorization from UCSD in compliance with institutional guidelines and procedures. Vero E6 (ATCC, CRL-1586) were maintained in Dulbecco’s modified eagle’s medium (DMEM) supplemented with 1% penicillin and streptomycin (P/S) and 10% fetal bovine serum (FBS) and cultured under 5% CO_2_ in a humidified incubator at 37^°^C.

### *In vitro* transcription of RNA

RNA was in vitro transcribed from 1 *μ*g of agarose gel-purified band corresponding to the intended size using HiScribe™ T7 high yield RNA synthesis kit (cat#E2040S, New England Biolabs Inc) followed by adding a 5’ cap using the Vaccina Capping System (cat#M2080S, New England Biolabs Inc.) and a poly-A tail using *E*.*coli* Poly(A) polymerase (cat#M0276S, New England Biolabs Inc.). Transcribed RNA was purified using phenol-chloroform extraction, followed by ethanol precipitation and resuspended in nuclease-free water.

### LNP formulation and characterization

RNA was packaged into LNPs using a NanoAssemblr microfluidic system (Precision Nanosystems, Vancouver, Canada) according to manufacturer’s instruction. Briefly, LNP formulations were prepared by injecting 12.5mM of lipid solution and 0.173*μ*g/*μ*l of RNA in formulation buffer at a flow rate of 12 ml/min. LNP suspension was immediately diluted in PBS (cat# 21-040-CM, Corning, Manassas, USA) followed by reconcentration of formulation by centrifuging at 2000 g in Amicon filters (30,000 MWCO, Amicon Ultra-15 Centrifugal filter unit cat# Z717185, Millipore Sigma, Burlington, USA). The LNP suspension was filtered through a 0.22-micron syringe filter, and LNPs were stored at 4^°^C until use. Free and total RNA concetrations were determined by Ribogreen assay (Quant-iT RiboGreen RNA, cat# R11490, Invitrogen, Carlsbad, USA). LNPs were lysed for 10 min at 37^°^C in 1% Triton X-100 to obtain total RNA concentration. Encapsulated RNA was calculated as ([total RNA]-[free RNA])/[total RNA] x 100). The size of LNPs was characterized by Dynamic Light Scattering in a DynaPro NanoStar Instrument (Wyatt Technology, Santa Barbara, USA) and analyzed with Dynamics 8.0 software (Wyatt Technology, Santa Barbara, USA). The LNPs were used within 5 days of making them.

### Transmission experiment

All Syrian golden hamsters (Hamster/Golden Syrian Hamster/Male/6-8 weeks old/Charles River/Strain Code 049) were approved by the Scripps Research Institute Institutional Animal Care and Use Committee/Protocol 20-0003) and experiments were carried out in accordance with recommendations. 8-week-old Syrian golden hamsters were intranasally infected with 10^6^ PFU of SARS-CoV-2 (USA-WA1/2020) or SARS-CoV-2 variant (B.1.617.2) in 100*μ*l of DMEM, as described (35). At 6-h post infection (for B.1.617.2 infected hamsters; 12-h post infection for USA-WA1/2020), hamsters were intranasally dosed with 100 *μ*l of LNP solution from either TIP RNA (n=5) or Ctrl RNA (n=5). At 36-h post infection, the source animals were co-caged with naïve animals (contact animals) for 8 hours, then all animals were caged individually starting 44-h post-infection. Nasal washes were collected for source animals on day 3, 4 and 5, and for contact animals on day 4, 5, and 6 followed by harvesting of lungs at day 5 for source and day 6 for contact animals.

### Plaque assay

Infectious virus was quantified by plaque assay on Vero E6 cells. Briefly, Vero E6 cells were plated as a confluent monolayer in 12-well plates 24 hours before performing the plaque assay. On the day of plaque assay, media was aspirated followed by washing cells with 2ml of PBS. Virus dilution was performed in modified DMEM media (DMEM, 2% FBS, L-glut, P/S), followed by adding 250 *μ*l of diluted virus to the confluent monolayer. The plates were incubated at 37^°^C for 1 hour with gentle rocking every 15 minutes. After one hour of incubation, 2ml of overlay media (1.2% Avicel in 1X MEM) was added to each well and transferred to incubator. At 3-days post infection, overlay media was gently aspirated, and the monolayer was washed with PBS, fixed with 10% formalin for 1 hour and stained with 0.1% crystal violet followed by washing with cell culture grade water. The plaques were enumerated and virus titer was calculated to PFU/ml or mg of tissue.

### RNA extraction and qRT-PCR titering of virus

At indicated time points, lung homogenate or nasal washes were lysed in TRIzol LS (cat# 10296010, Invitrogen), using 0.75ml TRIzol LS for 0.25ml sample volume. RNA was extracted using the Direct-zol RNA extraction kit (cat#R2070T, Zymo Research Inc.). RNA was DNAse treated using RNAse-free Dnase I (cat#EN0521, Thermofisher Scientific). 1 *μ*g of RNA was reverse transcribed using SuperScript II Reverse Transcriptase with oligo d(T) primers (cat#12574026, Thermofisher Scientific), and cDNA was analyzed by quantitative real-time polymerase chain reaction (qRT-PCR) analysis using SYBR green PCR master mix (cat#4309155, Thermofisher Scientific) with sequence specific primers. All the lung homogenate samples were normalized to beta-actin (***SI Appendix***, Table S1).

### Histopathology

Formalin-fixed lung from each animal was processed and paraffin embedded, and tissue sections were stained with hematoxylin and eosin (H&E) as described (36). Samples were imaged and analyzed using the Leica Aperio ImageScope software. Histopathological scoring was performed based on established algorithms (37-39) to devise a continuous numerical scale for determining the degree of pathogenicity in lung specimens for correlation with viral titer (PFU and qPCR) and inflammatory expression data. Specifically, the multiparametric quantitative scoring system analyzed a blinded histology image that considered the prevalence of pulmonary infiltrates, edema, macrophages, and septum widening, resulting in a composite histopathological score ranging from a minimum of 0 (indicating the absence of visual indications of pathogenicity in all scoring dimensions), to a maximum of 8 (indicating end-stage pathogenesis evidenced by overwhelming infiltrates, and/or edema, macrophage, and septum widening). Lungs with a score of 3 or lower were considered to be healthy lungs, with predominantly unobstructed alveolar capacity (37-39).

### Mathematical modeling and model inference

The following system of ordinary differential equations was used to model viral dynamics within individual hamsters:

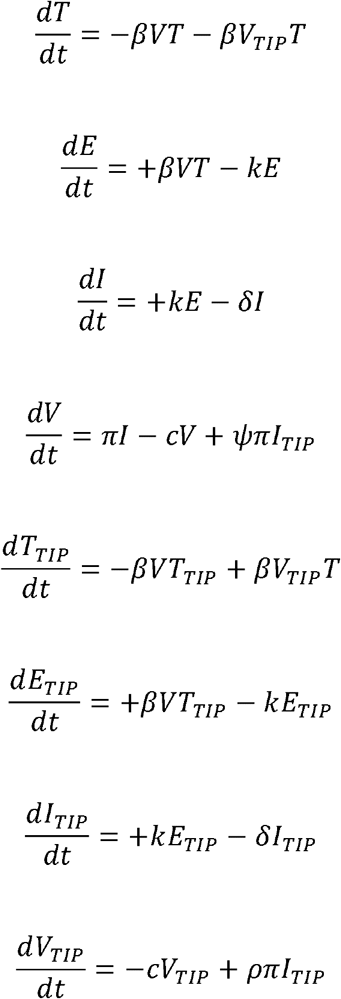

where T corresponds to naïve target cells, E corresponds to non-productively infected cells, I corresponds to productively infected cells, and V corresponds to SARS-CoV-2 viral load. These discrete states and their transitions are typical of viral dynamics models (30, 40, 41). We extend this model to have parallel TIP-carrying states: T_TIP_ corresponds to TIP-carrying target cells, E_TIP_ corresponds to TIP-carrying non-productively infected cells, I_TIP_ The parameter *β* is the infectivity, k is the rate of progression to productive infection, is the corresponds to TIP-carrying productively infected cells, and V_TIP_ corresponds to TIP load. death rate of infected cells, *c* is the TIP and virus clearance rate, *ρ* is the virus production rate, *ρ* is the TIP production rate relative to SARS-CoV-2, and *ψ* is the SARS-CoV-2 production rate in *I*_*TIP*_ cells relative to untreated infected cells. We assume *ρ* = 1.5 and *ψ* = 0.02 based on prior measurements (21).

For Ctrl-treated sources and contacts, all the TIP-related state variables were set to zero. For TIP-treated sources, a non-zero V_TIP,i_ was defined representing the amount of TIP administered at t=8h. For contacts of TIP-treated animals, we assumed no TIP transmission by setting all the TIP-related state variables to zero. We assumed there are 10^7^ target cells (T) as a rough estimate of the number of SARS-CoV-2 susceptible cells in the hamster respiratory system (42).

A two-stage approach was used to fit the model to the nasal wash plaque assay data. First, initial parameter estimates were generated by fitting only control-treated source hamsters within a nonlinear mixed effects model framework (Monolix version 2020R1. Antony, France: Lixoft SAS, 2020). At this stage, the rate of progression to productive infection, *k*, was fixed to 4 days^-1^ and the virion burst size π to 10 PFU mL^-1^ day^-1^ under the assumption that the single cell replication kinetics would not change substantially between hamsters. By fitting the control source dataset, we obtained representative parameter estimates for the population of control-treated source hamsters. For the next stage, these estimates were used to set priors for the Delayed-Rejection Adaptive-Metropolis variation of Markov Chain Monte Carlo (DRAM-MCMC) (43, 44). DRAM-MCMC was used to fit the nasal wash parameters and predicted viral dynamics. Gaussian priors were used on *β, δ, C*, log10 (*V*_*i*_), plaque assay data for each hamster separately, obtaining posterior distributions of model and for TIP-treated source hamsters, log10(V_*TIP,i*_). The viral inoculum and TIP dosage were sampled in logarithmic space. We assumed the prior was strongly informative for *β* (μ _*β*_ = 0.000031,*μ* _*β*_ = 0.00001)but uninformative for 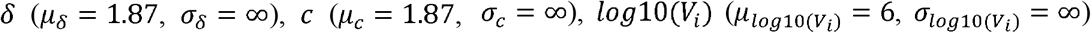 and 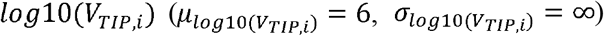. Narrowing the priors did not influence the quality of model fits. The sampling range for parameters was (*β*∈ [0,1], *δ*∈ [0,100], *c* ∈ [0,100], *log* 10 (*V*_*i*_) ∈ [1,7], *log* 10 (*V*_*TIP,i*_) ∈ [1,7]).

The following error function was minimized:

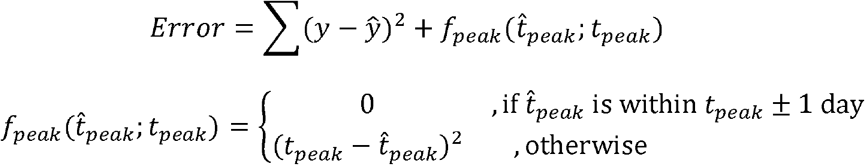

where ∑ (y-ŷ) ^2^ is the sum of squared errors from the data, and 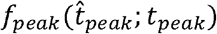 is a penalty function constraining the predicted timing of the peak viral load 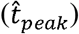 based on prior infection time course studies (28). Specifically, for source hamsters, we assume *t*_*peak*_ = 2, and for contact hamsters, we assume *t*_*peak*_ = 3. All simulated datapoints below the limit of detection (500 PFU/ml) were left-censored prior to calculating errors or variance defined by 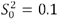 and N_0_= 20, which was empirically found to have a sufficient prediction intervals. We ran 10,000 steps of DRAM-MCMC using a prior for the error rejection frequency (∼40%) for sampling. The first 1,000 steps of DRAM-MCMC were discarded as burn-in, and 1,000 samples were drawn from the remaining chain to generate the posterior prediction intervals. The errors converged within this time frame (***SI Appendix*, Fig. S5a**) despite substantial uncertainty in individual parameters (***SI Appendix*, Fig. S5b**). The median of the prediction intervals (dark lines within shaded regions in **Fig. 4b, c**) were used to infer clearance time, integrated viral shedding, and peak viral shedding.

## Statistical analysis

Statistical differences were determined by using two-tailed unpaired Mann-Whitney U test, unless otherwise mentioned (GraphPad Prism). A p-value less than 0.05 was considered statistically significant: *<0.05, **<0.01, ***<0.001, ****<0.0001, ns: not significant.

## Data Availability

All unique reagents generated in this study are available from the Lead Contact with a completed Materials Transfer agreement.

