## Supplemental Figures 1-5 and Table 1 for "A single-administration therapeutic interfering particle reduces SARS-CoV-2 viral shedding and pathogenesis in hamsters"

### Supplemental Figure 1

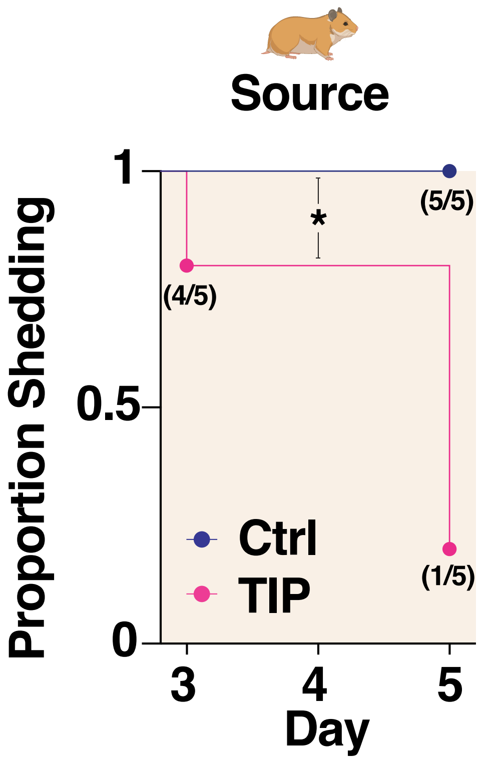

**Figure S1: Proportion of source hamsters shedding detectable virus quantified using the virus titer data in Fig. 1C and presented as a Kaplan-Meier plot.** *, $p<0.05$ by Log-rank (Mantel-Cox) test.

### Supplemental Figure 2

#### Source Ctrl

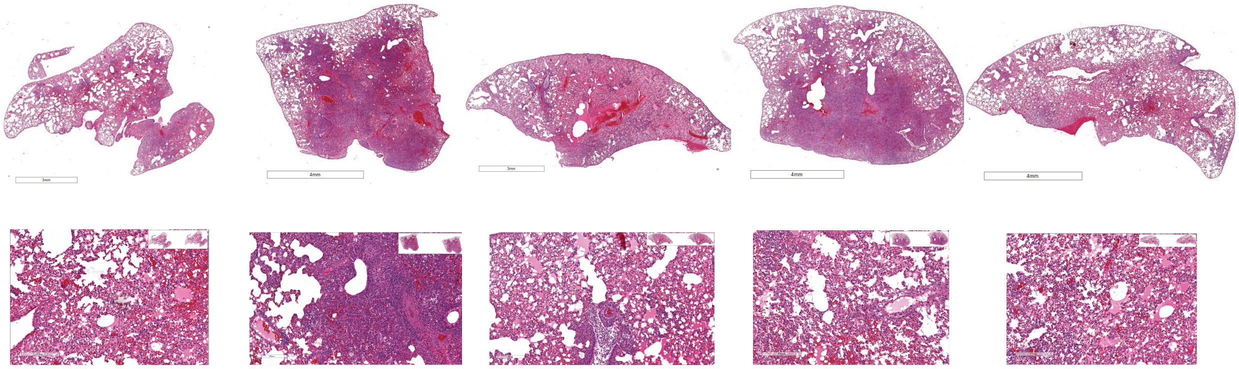

#### Source TIP

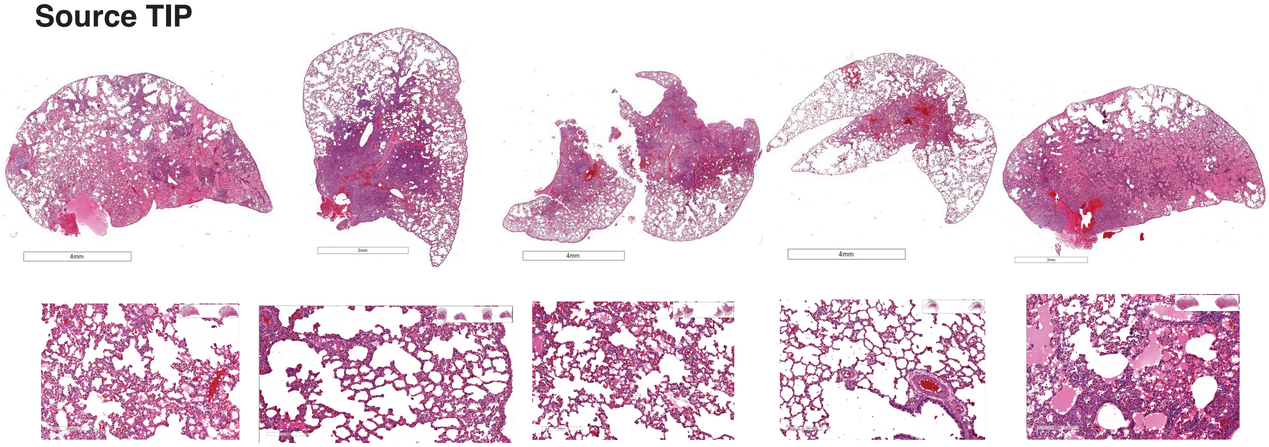

#### Contact Ctrl

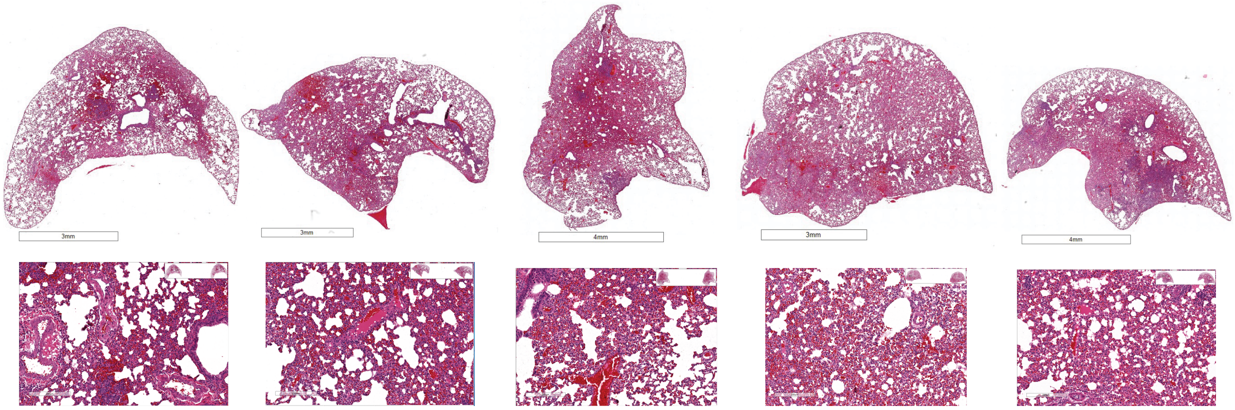

#### Contact TIP

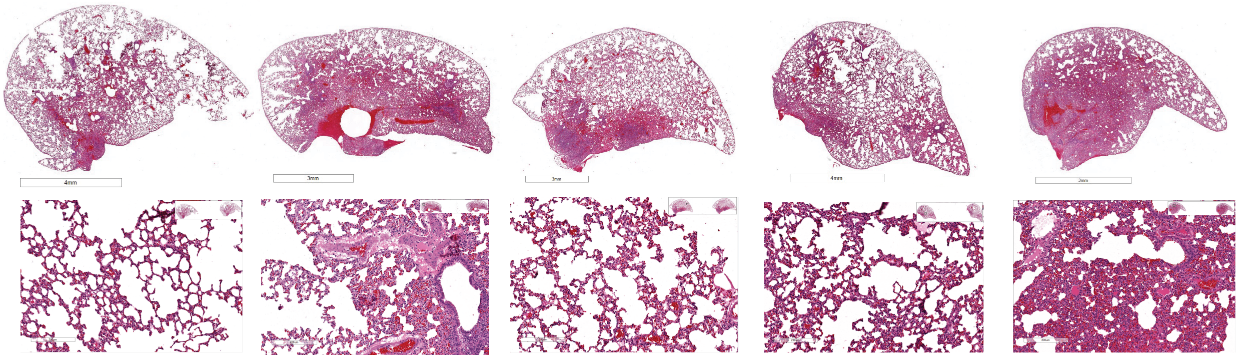

**Figure S2.: Histopathology imaging of Syrian golden hamster lungs from source animals infected with SARS-CoV-2 (B.1.617.2) followed by intranasal Ctrl- or TIP-LNP treatment and their respective contact animals.** Syrian golden hamsters (source) were intranasally infected with $10^6$ PFU of SARS-CoV-2 (B.1.617.2), at 6 h post-infection, intranasal administration of TIP LNPs (n=5) or Ctrl RNA LNPs (n=5) was performed. At 36 h post-infection, uninfected and untreated hamsters (contact) were brought in direct contact with the source hamsters for 8 hours. At 44 h post-infection, source and contact hamsters were caged alone, lungs of source and contact animals were harvested at day 5 and day 6 respectively, fixed with Zn-formalin and histopathology was performed. Sticked images were analyzed using Leica Aperio ImageScope software. For each animal a representative zoomed-in section to visualize histopathology are shown. Micrographs of brightfield imaging of H&E-stained lung sections from all animals are shown (*first row*: Ctrl RNA treated source (n=5), *second row*: TIP RNA treated source (n=5), *third row*: contact hamsters cohoused with Ctrl-treated hamsters (n=5), *fourth row:* contact hamsters cohoused with TIP-treated hamsters (n=5)).

#### Supplemental Figure 3

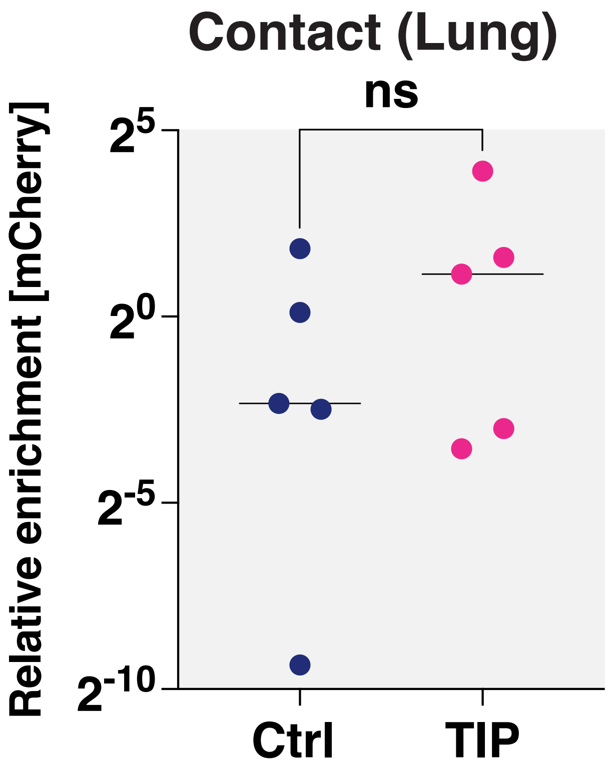

**Figure S3: No evidence for transmission of TIP to contacts from source animals infected with SARS-CoV-2 (B.1.617.2) followed by intranasal Ctrl- or TIP-LNP treatment.** Lungs from contacts (at day 6) of TIP-treated (n=5) or Ctrl-treated (n=5) animals were harvested, homogenized, and qRT-PCR was performed for mCherry (TIP marker) and normalized to beta-actin. ns, $p>0.05$.

### Supplemental Figure 4

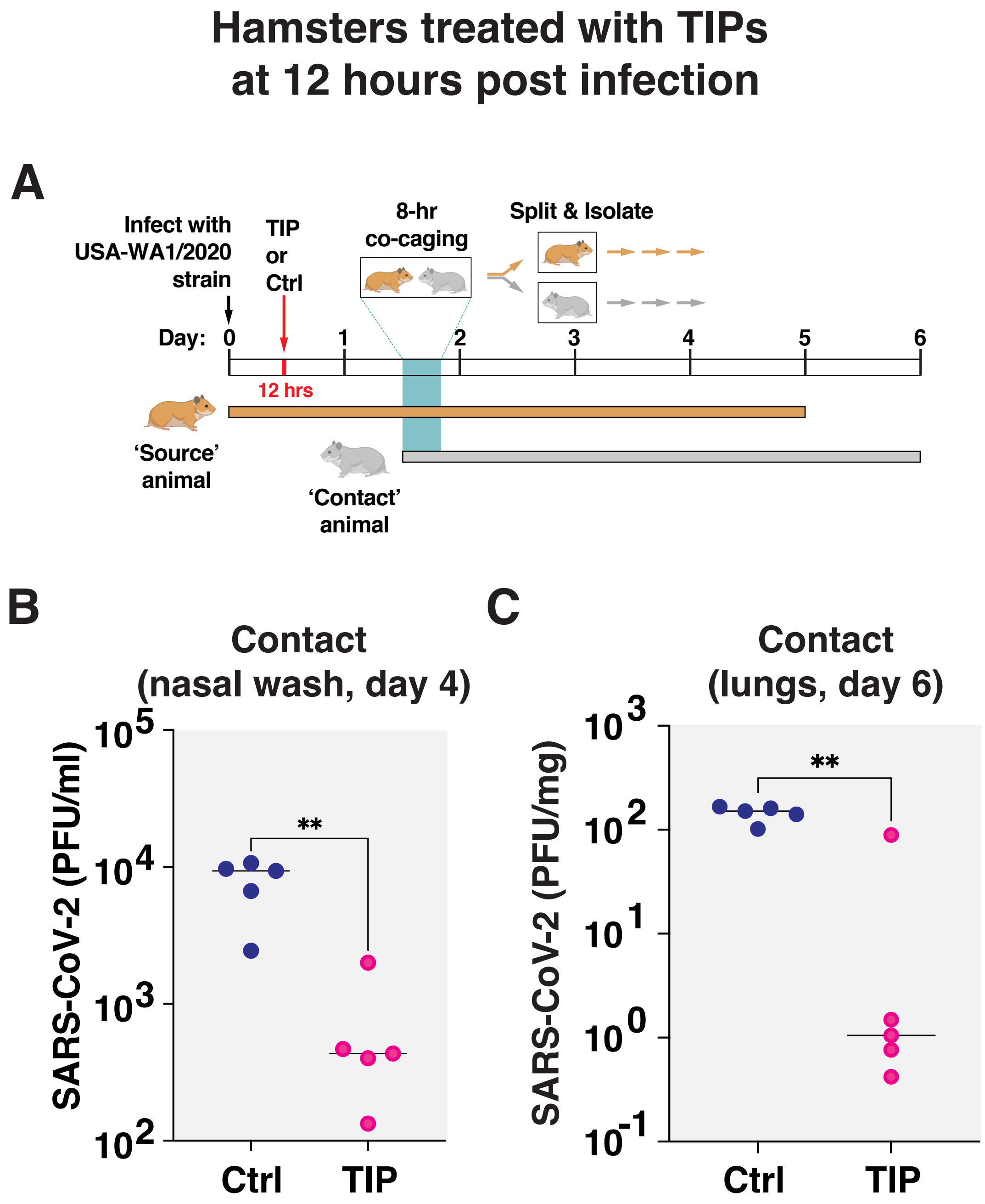

**Figure S4: TIP treatment post-infection (USA-WA1/2020) results in lower viral shedding and reduced viral load in the lungs of contact animals. (A)** Schematic of experimental design. Syrian golden hamsters (M/8-week-old) were intranasally infected with $10^6$ PFU of SARS-CoV-2 (USA-WA1/2020) in 100µl of DMEM, as described (29). At 12-h post, hamsters were intranasally administered with 100µl of LNP solution containing either TIP RNA (n=5) or Ctrl RNA (n=5), at 36-h post infection, the source animals were co-caged with naïve animals (contact animals) for 8 hours, and all animals were caged individually starting 44-h post-infection. Nasal washes were collected for contact animals on day 3 and 4 followed by harvesting of lungs at day 6 (5 days after co-caging). **(B)** Exposure to TIP-treated animals reduces virus shedding in contacts, as measured by plaque assay of nasal washes from day 4. **(C)** Exposure to TIP-treated animals reduces infectious viral load in lungs of contacts, as compared to exposure to Ctrl-treated animals. Lungs from contacts were harvested, homogenized, and SARS-CoV-2 virus titer was quantified by plaque assay.

### Supplemental Figure 5

**A**

#### Model error convergence

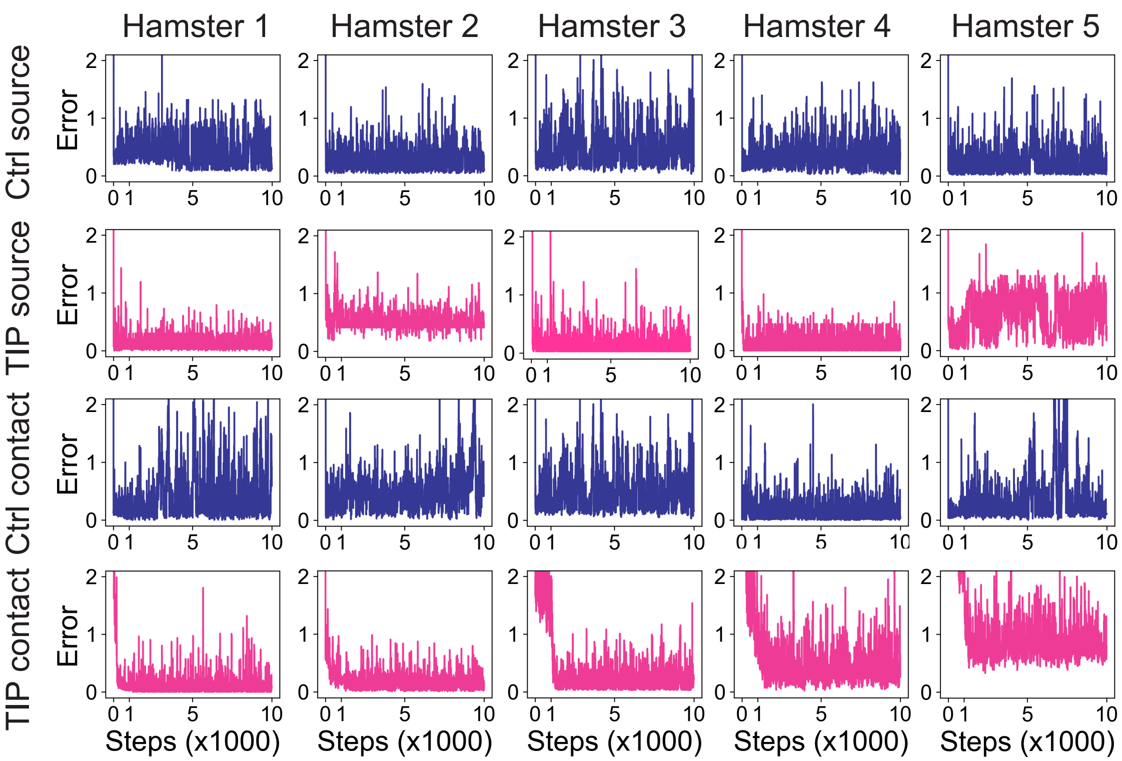


**B**

#### Model parameter distributions

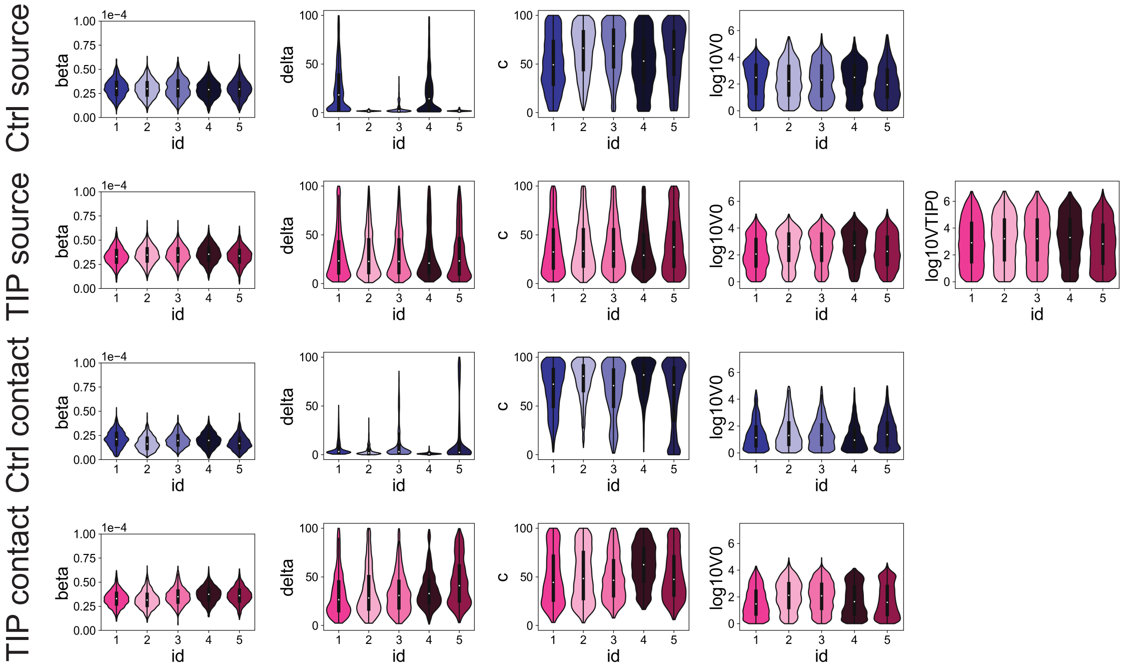

**Figure S5: Model fitting to source and contact hamster data. (A)** Diagnostic plots across the 10,000 DRAM-MCMC steps, exhibiting convergence of fitting error. The first 1,000 steps are discarded as burn-in, and the posterior distribution (parameters, and median prediction, and predicted credible intervals) are generated by sampling from the remaining 9,000 steps. **(B)** The posterior parameter distributions indicate that the values of individual parameters have large uncertainty. Parameter uncertainty is likely due to both structural non-identifiability, driven by parameters compensating for one another with no impact on the observed variable, and practical non-identifiability, driven by variation inherent in the data, and limited availability of time points.

**Table S1.** List and sequences of oligonucleotides used in the study.

| | | |
|---|---|---|
| 1 | N gene qpcr | fw: aaattttggggaccaggaac<br>rev:tggcacctgtgtaggtcaac |
| 2 | b-actin (M. auratus) qpcr | fw: actgccgcatcctcttcct<br>rev: tcgttgccaatggtgatgac |
| 3 | IL6 (M. auratus) qPCR | fw: ggtatgctaaggcacagcacact<br>rev cctgaaagcacttgaagaattcc |
| 4 | IL10 (M. auratus) qPCR | fw: gaaggaccagctggacaaca<br>rev:tggcaacccaagtaaccctta |
| 5 | TNF (M. auratus) qPCR | fw: ggagtggctgagccatcgt<br>rev:agctggttgtctttgagagacatg |
| 6 | IL1-beta (M. auratus) qPCR | fw: ggctgatgctcccattcg<br>rev: cacgaggcatttctgttgttca |
| 7 | CCL20 (M. auratus) qPCR | fw: agtcagtcagaagcaagcaact<br>rev: tgaagcggtgcatgatcc |
| 8 | IL4 (M. auratus) qPCR | fw: ccacggagaaagacctcatctg<br>rev: gggtcacctcatgttggaaataaa |
| 9 | IL10 (M. auratus) qPCR | fw: gttgccaaaccttatcagaaatga<br>rev: ttctggcccgtggttctct |
| 10 | IL2 (M. auratus) qPCR | fw: gtgcacccacttcaagctctaa<br>rev:aagctcctgtaagtccagcagtaac |
| 11 | CXCL10 (M. auratus) qPCR | fw: gccattcatccacagttgaca<br>rev: catggtgctgacagtggagtct |
| 12 | ISG15 (M. auratus) qPCR | fw: tctatgaggtccggctgaca<br>rev: gcactggggctttaggtcat |
| 13. | IFN- gamma (M. auratus) qPCR | fw: ggccatccagaggagcatag<br>rev: tttctccatgctgctgttgaa |
| 14. | TIPs PCR primers | t7 fw: taatacgactcactatagggattaaaggtt<br>rev: ttttttttttttttttgtcattctcctaagaagc |
| 15. | mCherry qPCR | fw: gaacggccacgagttcgaga<br>rev: cttggagccgtacatgaactgagg |
